# Water Stress Tolerance, Genomic Selection and Identification of Genomic Regions in a MAGIC Population of Eggplant

**DOI:** 10.1101/2025.08.18.670809

**Authors:** Martín Flores-Saavedra, Jon Bančič, Yuliza Huaman, Andrea Arrones, Oscar Vicente, Mariola Plazas, Santiago Vilanova, Pietro Gramazio, Jaime Prohens

## Abstract

Horticultural crops are increasingly affected by water stress due to climate change, making the development of stress-tolerant varieties an urgent need. In this study, we evaluated a set of 184 Multi-parent Advanced Generation Inter-Cross (MAGIC) eggplant lines under water stress conditions, consisting of irrigation at 30% of field capacity. After 21 days of stress, we assessed traits related to growth, water content, plant pigments, and proline content. The MAGIC population displayed a high variability for water stress tolerance, as transgressive lines over parental values were found for most of the traits. Key traits associated with water stress tolerance were identified, including increased root growth and elevated proline, and flavonoid content. Genome-Wide Association Study (GWAS) analysis allowed identification of three genomic regions associated with total dry weight, water content and flavonoids, traits that contribute significantly to stress tolerance. Additionally, a linear genomic selection index was constructed based on total dry weight, dry weight increase during the stress period, root dry weight, water content, and proline content to identify water stress-tolerant and susceptible lines. The model’s predictive ability was calculated, and the value of the selection index was predicted for 141 unevaluated MAGIC lines, with prediction accuracy for the index traits ranging from 0.11 to 0.53. This study presents a comprehensive analysis, identifying critical traits, genomic regions and lines with genetic potential, providing valuable information for future breeding programmes to improve water stress tolerance in eggplant.

## Introduction

Crops are frequently exposed to abiotic stresses such as high temperatures, drought, and salinity, which disrupt plant development and lead to significant yield losses [1,2]. Climate change has exacerbated drought conditions in many regions by increasing evapotranspiration and raising water demand, while simultaneously reducing precipitation and irrigation availability [3]. In horticultural systems, drought is expected not only to reduce yield but also to impact crop quality and increase susceptibility to pests [4].

Among horticultural crops, eggplant is of high economic importance, and its global production has steadily increased over the last 50 years [5,6]. Although water stress negatively affects eggplant (*Solanum melongena* L.) yield, it is relatively more tolerant than other *Solanum* vegetable crops [7], as eggplant can tolerate mild water stress without significant yield losses [8]. Physiologically, photosynthesis, transpiration and stomatal conductance of eggplants are directly affected by stress and decrease with reduced irrigation [9,10]. In addition, drought conditions induce oxidative stress in eggplant, though there is considerable genotypic variation, with more tolerant lines exhibiting enhanced antioxidant responses, including increased flavonoid accumulation under stress [11,12]. On the other hand, eggplant has been found to increase its proline content during stress periods, generating an osmoregulation that allows a lower reduction in photosynthesis due to water stress [13,14].

Considering the complexity and multifactorial nature of drought tolerance, an interdisciplinary approach is required, with plant breeding playing a crucial role [15]. Despite extensive research on abiotic stress tolerance mechanisms, breeding efforts in commercial crops remain primarily focused on optimal growing conditions [16]. While various drought tolerance mechanisms have been identified, incorporating them into breeding programmes remains challenging due to their complexity [17]. The development of Multi-parent Advanced Generation Inter-Cross (MAGIC) lines has significantly advanced plant breeding by facilitating the selection of recombinant lines with high potential for breeding programmes and enabling the identification of genomic regions and candidate genes [18,19]. MAGIC populations have demonstrated promising results in abiotic stress tolerance studies across various crops, including cotton [20], groundnut [21], bean [22], maize [23] and chickpea [24]. In eggplant, the only existing MAGIC population [25] has shown great potential for detecting genes and genomic regions of interest for breeding [26–28]. Interestingly, this population incorporates wild genetic diversity from *S. incanum* L., a drought-tolerant species known to harbour drought-related genes [29]. Additionally, the other seven *S. melongena* parental lines of the MAGIC population have displayed diverse biochemical and growth responses under water stress conditions [11], making this population particularly valuable for studying drought tolerance mechanisms and enhancing breeding strategies.

Given the polygenic nature and low heritability of certain traits, genomic selection has emerged as a powerful tool for accelerating genetic gains by integrating phenotypic and genomic information [30]. Genomic selection has been extensively applied to enhance yield, tolerance to biotic and abiotic stresses, and quality in cereals and industrial crops. However, its adoption in horticultural crops remains limited [31]. Genomic selection has been successfully used to predict the genetic value of complex traits in MAGIC populations [32,33], with moderate predictions also obtained in populations studied under water stress conditions [34].

This study aims to evaluate the variation in drought tolerance and related traits in a set of 184 genetically diverse eggplant MAGIC recombinant inbred lines and their parental lines under controlled water stress conditions. By integrating phenotypic and genomic data, we aim to identify recombinant MAGIC lines with improved performance, detect genomic regions associated with key drought-related traits, and apply genomic selection to more accurately select for drought tolerance in eggplant.

Furthermore, we assess the predictive ability of these models and use them to predict drought tolerance in lines that were not evaluated, supporting the use of genomic selection as a tool for future drought-tolerance improvement in eggplant.

## Results

### Trait analysis

MAGIC lines showed higher average values than the parents for most evaluated traits under the water stress conditions used, except for Total water content (WC), anthocyanin index (Anth), Aerial WC, Stem WC, Leaf number growth, Leaf WC, and Root dry weight (DW) growth (Table 1). Moreover, the MAGIC lines exhibited a broader range of variation than the parents for nearly all traits, with transgressive individuals exceeding both lower and upper parental values except for Total WC and Stem WC, where no lines fell below the lower parental values (Table 1).

**Table 1.** Mean and range for parents and MAGIC lines. Only the MAGIC lines were used to estimate phenotypic coefficient of variation (pCV), genotypic coefficient of variation (gCV), broad-sense heritability (H²), and narrow-sense heritability (h²) for each trait.

| Trait<sup>a</sup> | Parents | | MAGIC lines | | pCV | gCV | $H^2$ | $h^2$ |
| --- | --- | --- | --- | --- | --- | --- | --- | --- |
|  | Mean | Range | Mean | Range |  |  |  |  |
| Stem Length (cm) | 12.35 | 10.06-15.37 | 13.77 | 7.07-26.40 | 23.42 | 21.34 | 0.84 | 0.84 |
| Stem DW (g) | 0.71 | 0.52-0.88 | 0.88 | 0.28-1.65 | 27.06 | 21.12 | 0.77 | 0.60 |
| Anth | 0.35 | 0.32-0.39 | 0.34 | 0.27-0.43 | 7.89 | 5.20 | 0.71 | 0.46 |
| Flav | 1.36 | 1.3-1.47 | 1.37 | 1.12-1.68 | 7.89 | 5.77 | 0.69 | 0.46 |
| Total WC (%) | 83.75 | 78.62-85.98 | 83.74 | 78.85-86.57 | 1.52 | 1.08 | 0.68 | 0.43 |
| Leaf WC (%) | 85.23 | 80.25-87.85 | 85.14 | 79.09-87.96 | 1.54 | 0.97 | 0.65 | 0.11 |
| Stem WC (%) | 78.28 | 68.68-81.53 | 77.65 | 70.40-84.76 | 3.00 | 2.32 | 0.63 | 0.63 |
| Leaf Number | 4.98 | 3.33-6.30 | 5.21 | 2.97-9.97 | 24.35 | 18.70 | 0.63 | 0.63 |
| Aerial WC (%) | 83.85 | 77.90-86.44 | 83.72 | 76.36-86.78 | 1.97 | 1.19 | 0.61 | 0.27 |
| Aerial/Root ratio | 2.63 | 1.69-4.09 | 3.66 | 1.53-8.41 | 24.27 | 17.69 | 0.58 | 0.58 |
| Chl | 39.60 | 34.63-43.06 | 42.87 | 31.77-54.21 | 9.40 | 6.78 | 0.57 | 0.57 |
| Leaf number growth | -0.49 | -1.31-0.33 | -0.96 | -3.78-2.22 | 10.61 | 4.84 | 0.57 | 0.10 |
| Stem length growth (cm) | 3.49 | 2.84-4.73 | 3.65 | 0.11-8.11 | 42.09 | 30.28 | 0.57 | 0.57 |
| NBI | 28.86 | 23.43-32.66 | 31.64 | 18.38-41.91 | 9.21 | 6.61 | 0.57 | 0.57 |
| Stem DW growth (g) | 0.51 | 0.37-0.57 | 0.60 | 0.17-1.05 | 30.59 | 21.70 | 0.56 | 0.56 |
| Root WC (%) | 82.74 | 80.06-84.65 | 83.18 | 78.01-88.01 | 2.23 | 1.50 | 0.51 | 0.51 |
| Aerial DW (g) | 2.86 | 2.18-3.67 | 3.68 | 1.44-5.82 | 20.24 | 13.17 | 0.49 | 0.49 |
| Leaf DW (g) | 2.15 | 1.67-2.79 | 2.80 | 1.16-5.09 | 21.25 | 13.76 | 0.48 | 0.48 |
| Total DW (g) | 3.91 | 3.13-4.79 | 4.75 | 2.00-7.11 | 19.25 | 11.71 | 0.44 | 0.44 |
| Root DW (g) | 1.05 | 0.82-1.17 | 1.08 | 0.57-1.70 | 24.41 | 14.46 | 0.42 | 0.42 |
| Root DW growth (g) | 0.80 | 0.68-0.98 | 0.78 | 0.40-1.33 | 29.58 | 14.79 | 0.33 | 0.33 |
| Pro ($\mu\text{mol g}^{-1} \text{DW}$) | 91.18 | 57.93-111.49 | 108.16 | 40.67-173.63 | 25.80 | 12.52 | 0.31 | 0.31 |
| Aerial DW growth (g) | 1.68 | 1.25-2.14 | 2.12 | 0.68-3.66 | 26.45 | 11.67 | 0.28 | 0.28 |
| Total DW growth (g) | 2.52 | 1.97-2.90 | 2.91 | 1.24-4.57 | 23.83 | 8.76 | 0.21 | 0.21 |
| Leaf DW growth (g) | 1.17 | 0.86-1.60 | 1.52 | 0.48-2.96 | 29.15 | 9.18 | 0.16 | 0.16 |

<sup>a</sup>WC, water content; DW, dry weight; Chl, chlorophyll index; Flav, flavonol index; Anth, anthocyanin index; NBI, nitrogen balance index; Pro, proline. Traits ending in “growth” refer to growth during the period of water stress.

The phenotypic (pCV) and genotypic (gCV) coefficients of variation varied notably among traits and were strongly correlated (r = 0.90). The traits exhibiting the highest phenotypic variation were primarily related to growth, such as Stem length growth, Stem DW growth, Stem DW, Root DW growth, and Leaf DW growth (Table 1). Similarly, the traits with the highest genotypic variation were those related to stem characteristics, including Stem length growth, Stem DW growth, Stem length, and Stem DW, followed by Leaf number and Aerial/Root ratio. On the other hand, the traits with the lowest phenotypic and genotypic variation were those related to water content, including Total WC, Aerial WC, Stem WC, Leaf WC, and Root WC, followed by pigment-related indices like chlorophyll index (Chl), flavanol index, (Flav), Anth, and nitrogen balance index (NBI) (Table 1).

The evaluated traits displayed a wide range of heritability values. Some traits, such as Stem Length, Stem DW and Anth, exhibited high H² values (>0.70). In contrast, traits related to growth during the stress period, including Aerial DW growth, Total DW growth, Leaf DW growth, had H² values below 0.30, likely due to higher experimental error associated with these measurements (Table 1). Values of h² were equal to H² for 18 traits due to the absence of residual additive variance (σ_n_ ^2^) (Figure 1). Among the traits with the highest h² values (greater than 0.60), we found Stem length, stem WC, Leaf Number and Stem DW. The traits with the lowest h² values were consistent with those having low H² values (Table 1).

**Figure 1.**
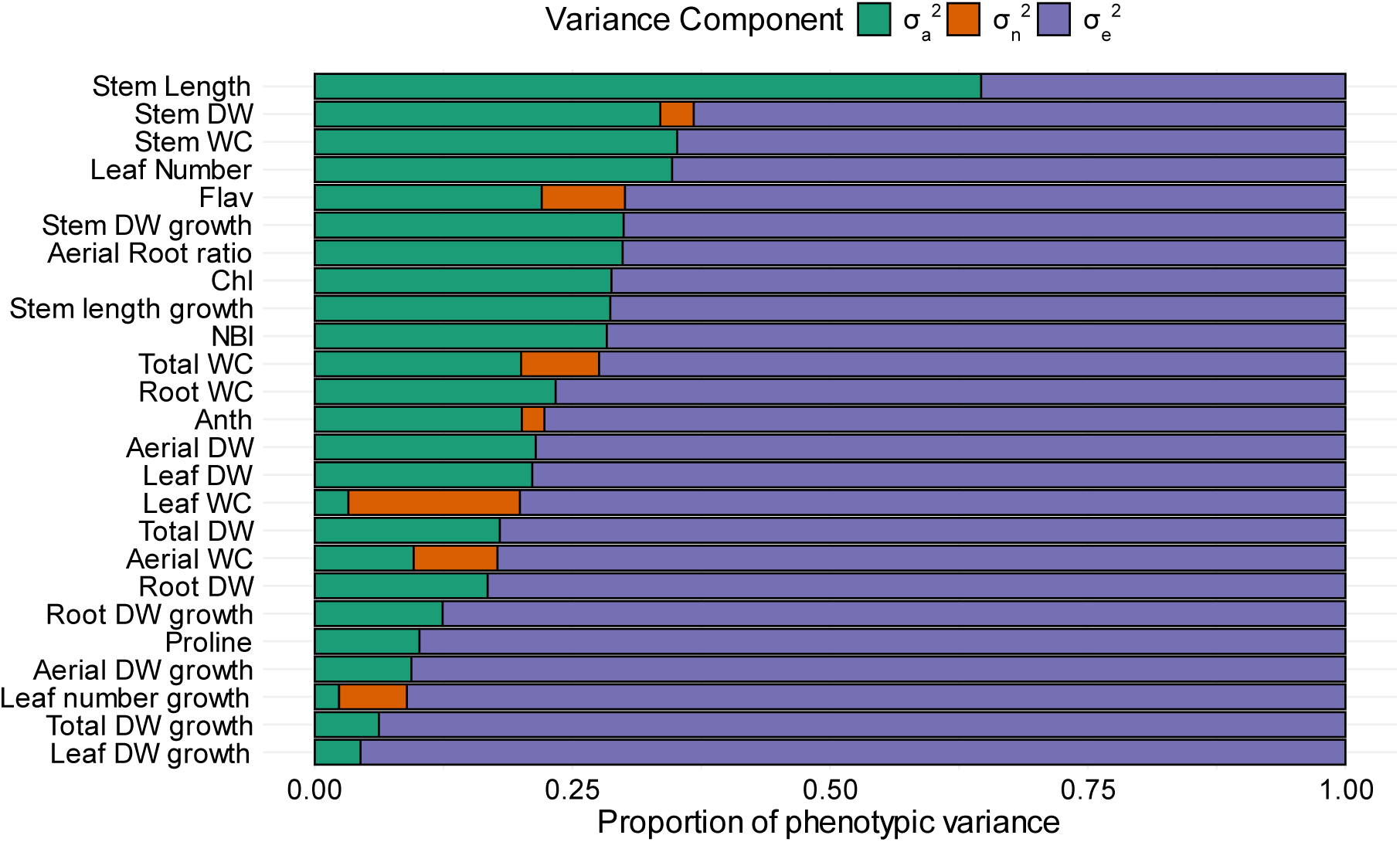
Variance components for each character assessed, where σ_a_^2^ is the additive variance, σ ^2^ is residual genetic variance and σ_e_^2^ is the residual variance. Traits are ordered according to decreasing values of Ve. WC, water content; DW, dry weight; Chl, chlorophyll index; Flav, flavonol index; Anth, anthocyanin index; NBI, nitrogen balance index. Traits ending in “growth” refer to growth during the period of water stress.

Seven traits exhibited σ ^2^ values (epistatic and dominance variance) greater than zero. Among these, the traits with the highest proportion of σ_n_ ^2^ to total phenotypic variance were Stem DW, Flav, Total WC, Anth, Leaf WC, Aerial WC, and Leaf number growth (Figure 1). The traits with the highest proportion of additive variance (σ_a_^2^) were Stem Length, Stem WC, Leaf Number, and Stem DW. On the other hand, traits associated with growth during the stress period, such as Leaf Number growth, Leaf DW growth, Total DW growth, Aerial DW growth, and Leaf WC had the lowest σ_a_^2^ values (Figure 1).

### Multivariate analysis

The correlations between total genetic values predicted from linear mixed model (Eq. 2) are shown in Figure 2 and Figure S1. Biomass traits (DW and DW growth) were highly correlated with each other, as were WC traits, but biomass traits tended to be negatively correlated with WC traits, with WC being lower when growth was higher (Figure 2; Fig. S1). Pigment-related traits, Chl, Flav were negatively correlated with WC traits, while Anth was positively correlated with WC. As for the correlation of pigments with biomass, Chl was positively correlated with Total, Aerial and Leaf DW, Flav was positively correlated with Total, Aerial, Leaf and Root DW, and Anth was negatively correlated with Total, Aerial and Leaf DW. Stem growth-related traits were positively correlated with aerial biomass and Stem WC and negatively correlated with Flav and root biomass. Leaf number traits were positively correlated with WC and negatively correlated with Total, Aerial and Leaf DW (Figure 2; Fig. S1). Meanwhile, Pro correlated positively with DW parameters related to Total Aerial and Leaf, and negatively with Root WC.

**Figure 2.**
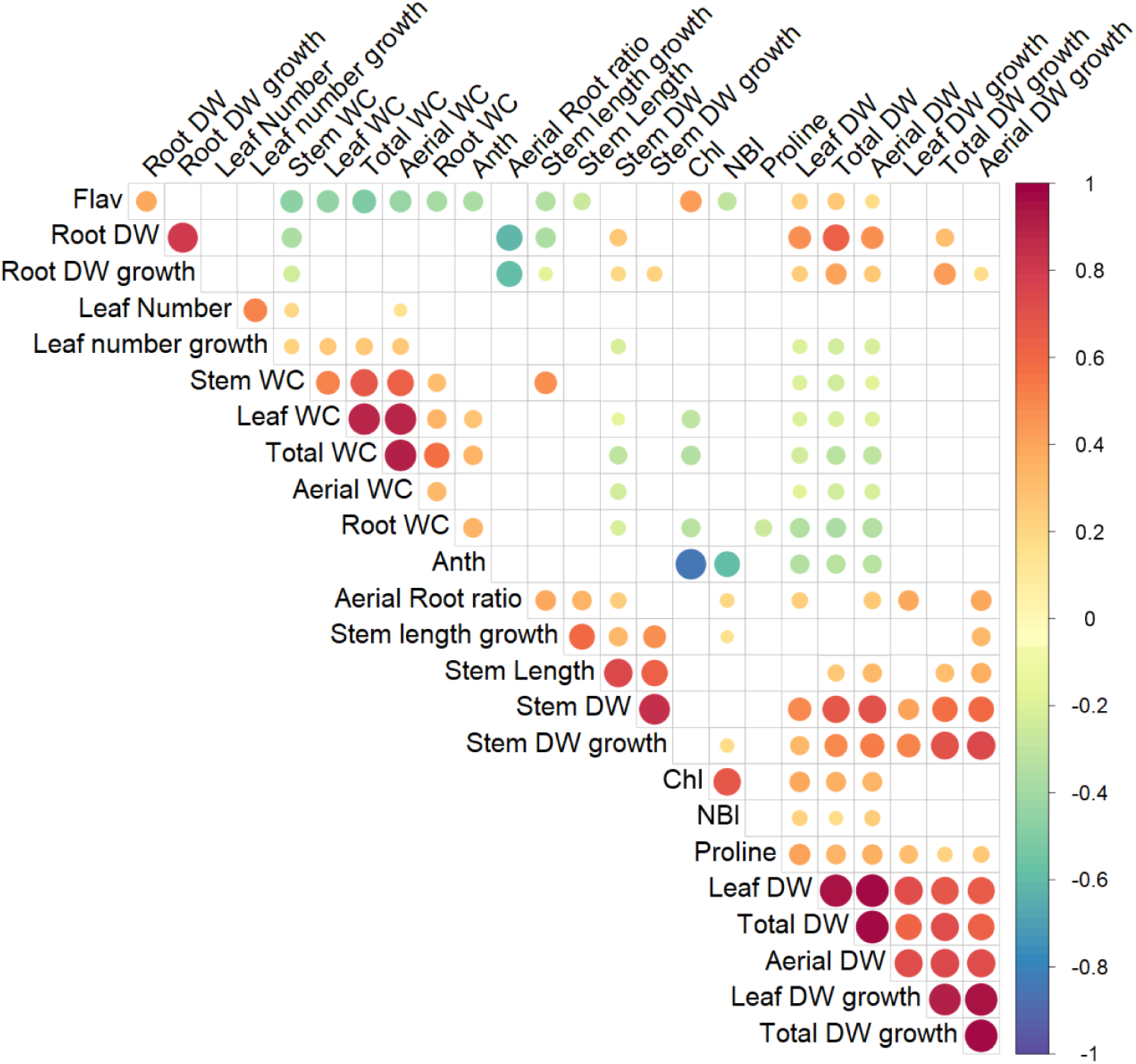
Genetic correlation matrix between traits in the subset of 184 eggplant MAGIC lines evaluated under water stress conditions. Only statistically significant correlations (*p* < 0.05) are shown. WC, water content; DW, dry weight; Chl, chlorophyll index; Flav, flavonol index; Anth, anthocyanin index; NBI, nitrogen balance index. Traits ending in “growth” refer to growth during the period of water stress.

The first two principal components of the principal component analysis (PCA), based on total genetic values predicted from the linear mixed model (Eq. 2), account for 48.4% of the observed variation, with the first principal component (PC1) and the second principal component (PC2) explaining 29.7% and 18.7%, respectively, of the total variation (Figure 3). In the loading plot, Total DW, Aerial DW, Stem DW, Leaf DW, Root DW, Total DW growth, Aerial DW growth and Leaf DW growth showed a highly negative relationship (< –0.5) with PC1, whereas Total WC displayed a highly positive relationship (> 0.5) with PC1. On the other hand, Total WC, Aerial WC, Stem WC, Leaf WC, Stem Length, Stem Length growth and Aerial/Root ratio showed high negative correlations (< –0.5) with PC2, whereas Flav displayed a high positive relationship (> 0.5) with PC2. In the score plot, all parents had positive values for PC1 and a wide range of values for PC2, while MAGIC lines were scattered over a broad area in the PC graph. It also shows that the MAGIC lines are mostly on the left side of the graph, unlike the parents (Figure 3).

**Figure 3.**
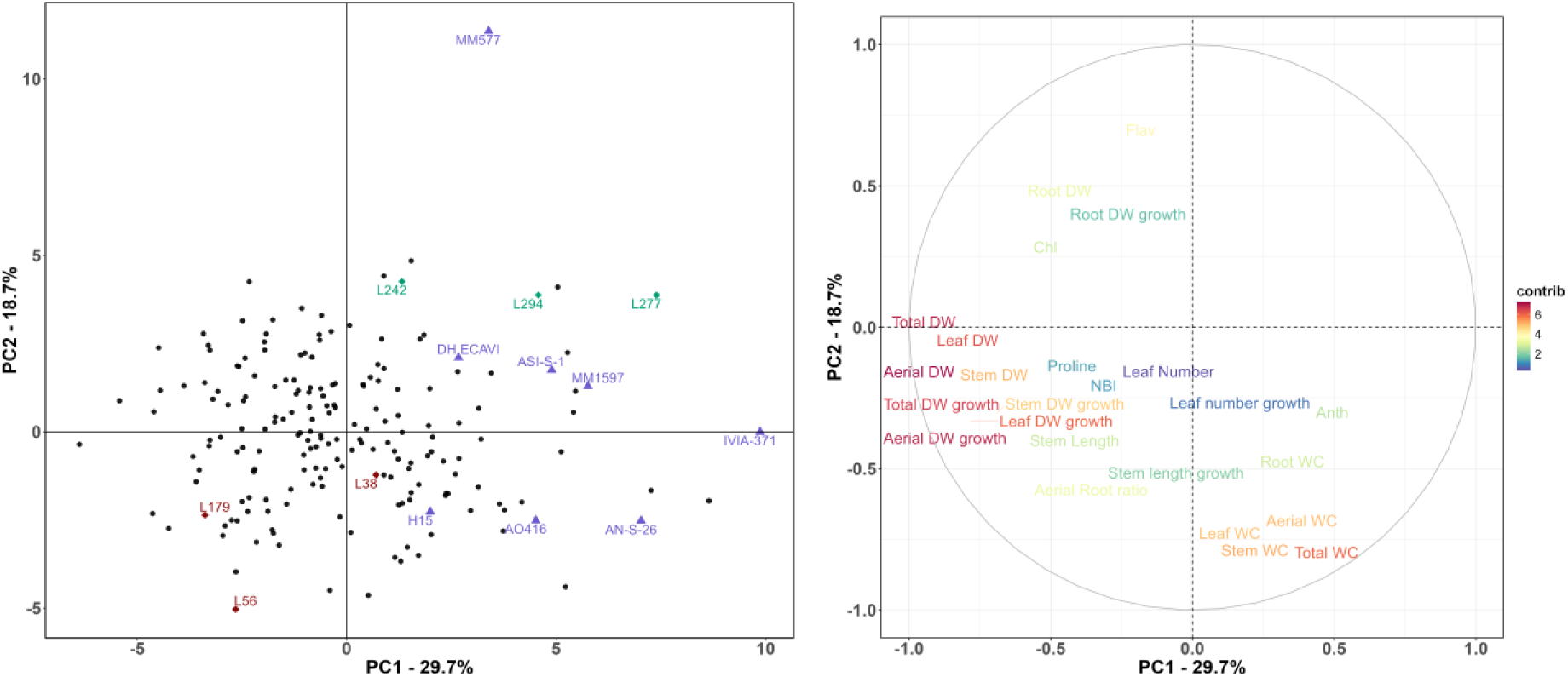
Score plot (left) and loading plot (right) of the principal component analysis (PCA) for the parents and the subset of 184 MAGIC lines, based on the first two principal components. The three most tolerant (L38, L56 and L179) and susceptible (L242, L277 and L294) lines, according to the Smith-Hazel selection indexes, are shown in red and green, respectively, and the parents are represented as purple triangles.

### Genome-wide association study and candidate genes

The results from the genome-wide association study (GWAS) analysis identified genomic regions significantly associated with three traits. These genomic regions were characterised by a region of significant SNPs with a clear upward trend in the areas (Fig. S2). A genomic region associated with the Total DW trait was found on chromosome 12, while two significant ones in different regions of chromosome 11 were associated with Total WC and Flav (Table 2).

**Table 2.** Genomic regions identified by GWAS, including –log10 (*p*), percentage of variance explained (R²), and minor allele frequency (MAF) for the most significant marker.

| Trait<sup>a</sup> | Chromosome | Position | $-\log_{10}(p)$ | $R^2$ | MAF |
| --- | --- | --- | --- | --- | --- |
| Total DW | 12 | 93447106-93874212 | 6.87 | 0.16 | 0.12 |
| Total WC | 11 | 3212497-3652760 | 7.34 | 0.16 | 0.11 |
| Flav | 11 | 60315426-62187141 | 7.10 | 0.17 | 0.15 |

<sup>a</sup>WC, water content; DW, dry weight; Flav, flavonol index.

Candidate genes were searched within a 1Mb region surrounding the genomic region for each trait. From a total of 31 genes in these regions for Total DW, 41 genes for Total WC, and 45 genes for Flav, three, five, and one candidate genes were identified as candidate genes for each trait, respectively.

The function of these candidate genes has been reported to be functionally related to the trait under water stress (Table 3).

**Table 3.** Candidate genes associated with the most significant marker for each trait in the GWAS analysis.

| Related trait<sup>a</sup> | Gene ID | Functional Annotation | References |
| --- | --- | --- | --- |
| Total DW | SMEL_012g394910.1 | Dof zinc finger protein DOF2.1 | Jin et al., 2024 |
| Total WC | SMEL_011g363040.1 | Inositol transporter INT4 | Zhou et al., 2022; Yildizli et al., 2018 |
|  | SMEL_011g363100.1 | Cysteine proteinase inhibitor Os03g0210100 | Zhang et al., 2008 |
|  | SMEL_011g363370.1 | Ubiquitin ligase regulating light response COP1 | Moazzam-Jazi et al., 2018 |
|  | SMEL_011g363380.1 | Tyrosine phosphatase involved in cellular signaling DSP1 | Liu et al., 2012 |
| Flav | SMEL_011g373070.1 | Transcription factor bHLH75 | Wang et al., 2016; Waseem et al., 2019 |

<sup>a</sup>WC, water content; DW, dry weight; Flav, flavonol index.

### Genomic selection

The linear genomic selection index enabled the identification of MAGIC lines exhibiting high growth under water stress conditions, along with elevated Pro contents and Total WC levels and moderate Flav contents (Figure 3). According to the index constructed, the most tolerant line to water stress was L56, followed by L38 and L179, while the most susceptible lines were L277, L242, and L294. The parents showed mean index values spread among the lines, with H15 and AN-S-26 being the parental with the highest values and ASI-S-1 the one with the lowest value (Figure 4). The lines selected as susceptible and tolerant also clustered differently in the PCA, with the susceptible lines at the top of the score plot and the tolerant lines at the bottom (Figure 3).

**Figure 4.**
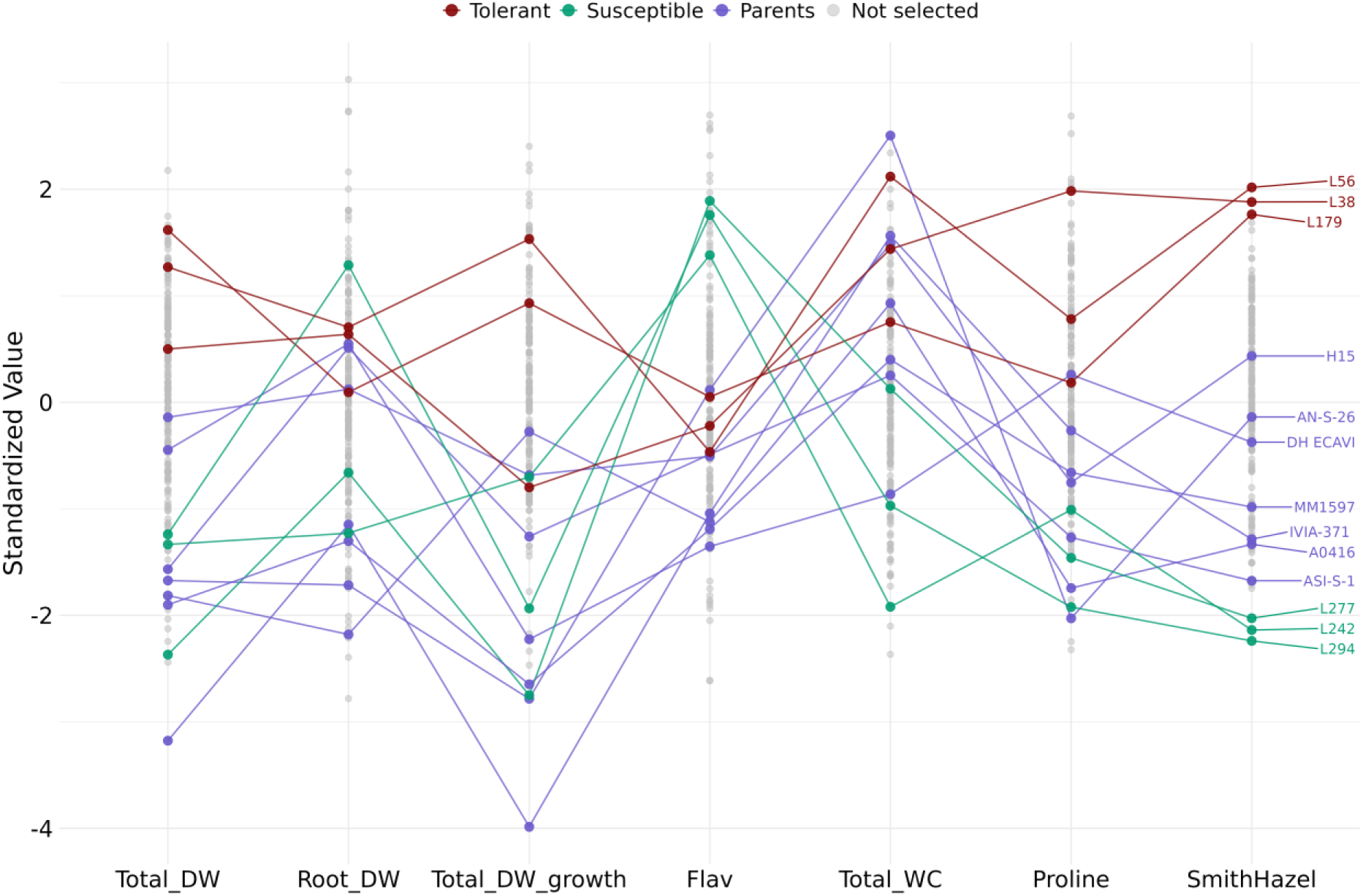
Linear genomic selection index constructed with six traits related to drought stress: Total DW, Total DW growth, Root DW, Flav, Total WC, and proline. The three lines with the highest index are labelled as tolerant (L56, L38, L179), while the three lines with the lowest index are labelled as susceptible (L277, L294, L213). Parentals (*S. melongena* only) are shown in purple. WC, water content; DW, dry weight; Flav, flavonol index.

The predictive ability of the model was calculated for each trait evaluated by the leave-one-out (LOO) and 5-fold methods; both methods were consistent in their results, with a correlation of 0.94, displaying similar predictive abilities for each trait (Table 4). Medium prediction accuracy (greater than 0.4) was observed for traits such as Leaf DW, Root WC, Chl, NBI, Stem DW growth, and ProIn contrast, the relationship between predicted and observed values was low for traits like Leaf WC, Flav, Leaf number and Leaf number growth, with prediction accuracy values of zero or even negative (Table 4).

**Table 4.**
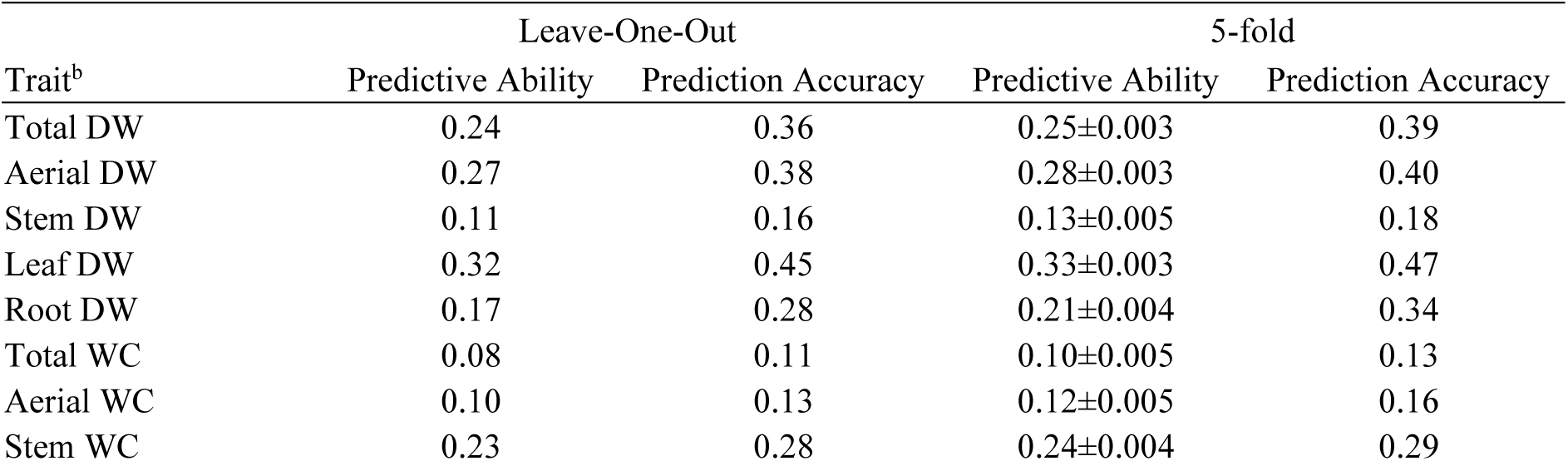
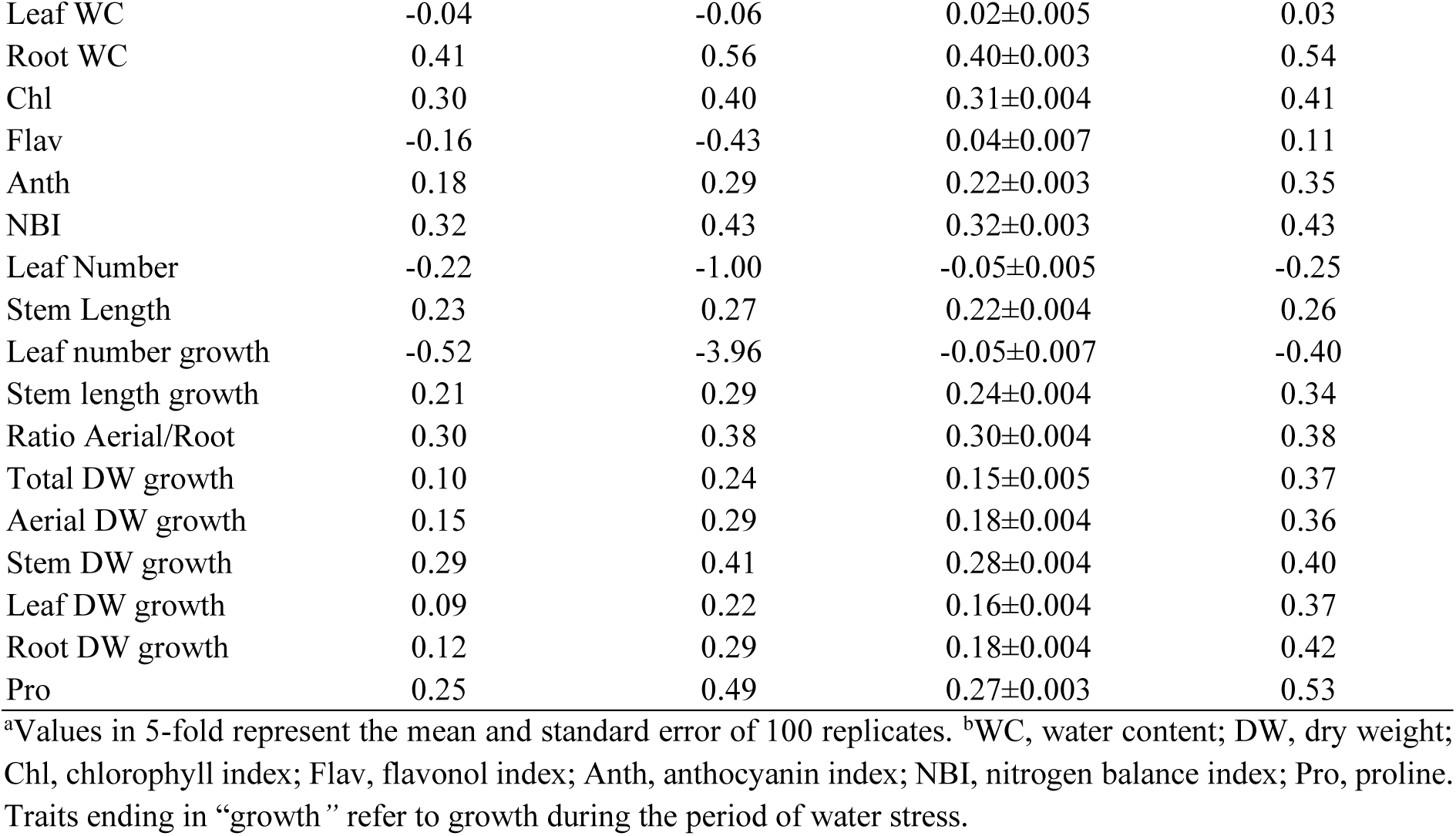
Predictive ability and prediction accuracy from cross-validation using the leave-one-out and 5-fold methodologies for each trait in the MAGIC population.

The selection model used predicted the 141 non-phenotyped MAGIC lines, calculated the GEBVs for each trait and calculated the selection index. In this way, the most tolerant lines (L323, L59 and L13) and the most susceptible lines (L234, L57 and L109) among the MAGIC lines not tested were predicted (Figure 5). The traits included in the index were Total DW, Root DW, Total DW growth, Flav, Total WC, and Pro, with prediction accuracies using the leave-one-out/5-fold cross-validation methods of 0.39/0.36, 0.34/0.28, 0.37/0.24, 0.11/-0.43, 0.13/0.11, and 0.53/0.49, respectively (Table 4).

**Figure 5.**
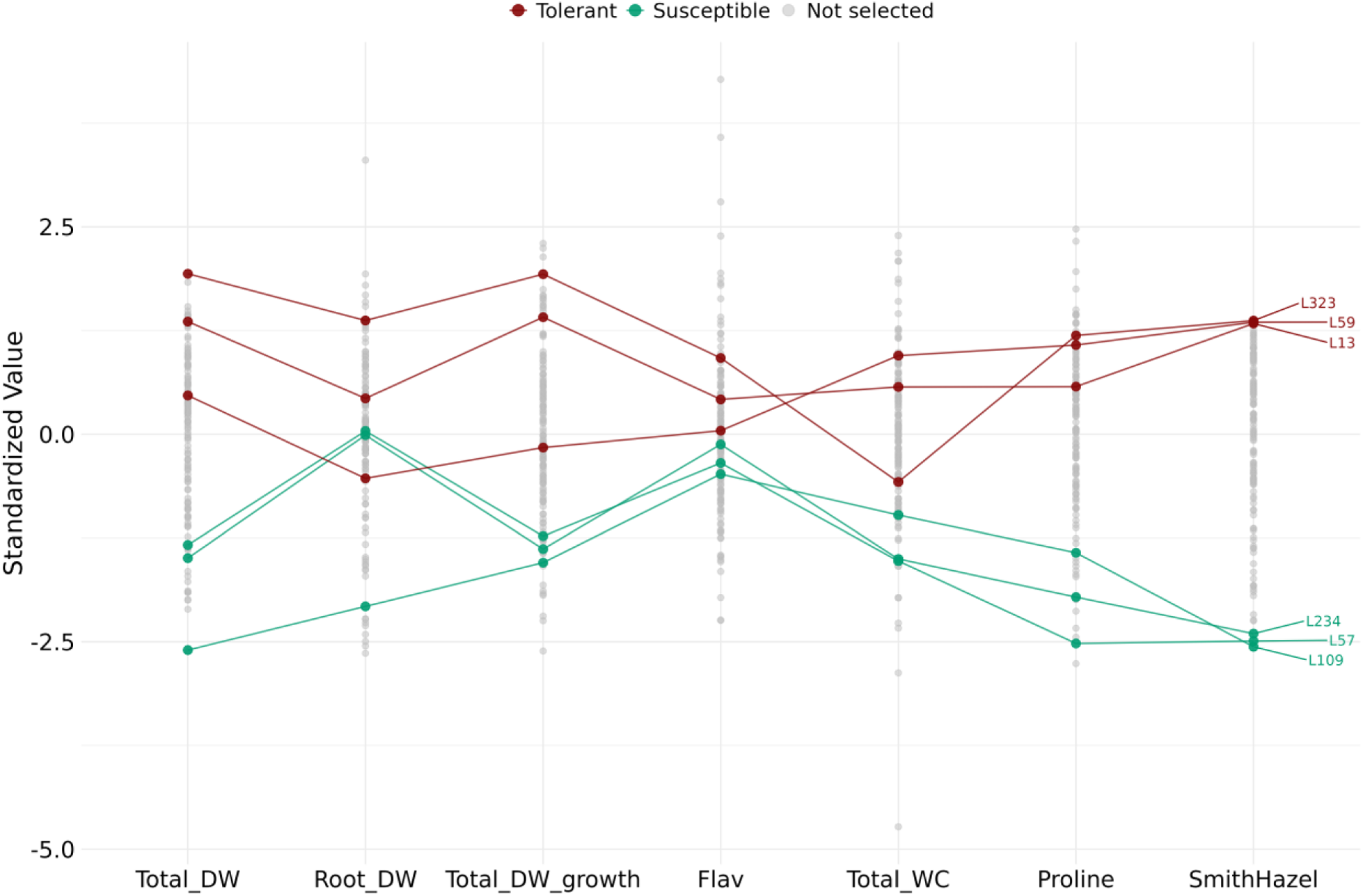
Linear genomic selection index based on genomic estimated breeding values (GEBVs) of six traits related to drought stress: Total DW, Root DW, Total DW growth, Flav and Total WC. This is for the genetic prediction of the MAGIC lines not included in the phenotyping experiment. The three lines with the highest index are labelled as tolerant (L323, L59, L13), while the three lines with the lowest index are labelled as susceptible (L109, L57, L234). WC, water content; DW, dry weight; Flav, flavonol index.

## Discussion

This study presents a comprehensive analysis of the mechanisms of water stress tolerance in the first MAGIC population of eggplant, focusing on three main aspects: phenotyping under stress conditions, identification of candidate genes through GWAS, and application of linear genomic selection index for improved drought tolerance. Together, these provide an innovative approach to understanding the tolerance mechanisms, genetic basis and selection of a drought-tolerant ideotype in eggplant. The results provide valuable information for future breeding programmes aimed at improving water stress resistance in this crop.

Plants are significantly affected by suboptimal water availability, and in the case of eggplant, water stress results in clear plant responses, such as leaf loss [35,36] and increased Pro content [11,37]. Progenies from crosses among genetically distinct parents often show transgressive inheritance in both morphological and physiological traits [38]; in this sense, the transgressive values observed in the eggplant recombinant MAGIC population for most traits suggest a wide diversity of responses to water stress, making it a valuable resource for plant selection and the identification of traits associated with increased tolerance to water stress.

Abiotic stress conditions often lead to reduced growth and biomass accumulation, primarily due to decreased energy supply from photosynthesis and lower water potential [39]. In this context, the ability to maintain growth under stress is considered an indicator of tolerance to reduced irrigation conditions. Among plant organs, some eggplant MAGIC lines exhibited significantly higher stem growth under water stress compared to the parental lines, with stem-related traits showing high heritability. However, stem growth was not found to be involved in eggplant drought tolerance [12]. Although leaf growth is potentially beneficial due to increased photosynthetic capacity, it also increases the transpiration surface area of the plant, making it more susceptible to dehydration [40]. In eggplant, it has been observed that lines with higher tolerance do not necessarily exhibit higher aerial biomass but rather higher root biomass [9]. Root growth is also recognised as an important trait in other *Solanum* crops [41,42], and in our study, we found lines with up to 45.3% and 35.7% higher Root DW and Root DW growth, respectively, compared to the best parent.

The negative relationship between growth and water content is logical, as bigger plants require more water for metabolic processes. However, growth is highly dependent on cell turgor [43]. Therefore, selecting for drought-tolerant plants requires identifying those with high growth potential that do not lose cell turgor, thus ensuring stable growth over time. Osmotic adjustment is an important tolerance mechanism that helps plants maintain cell turgor, and Pro is a key solute in this process [44]. Although Pro did not show a significant correlation with water content in our MAGIC population, it was positively correlated with biomass. Furthermore, Pro has been implicated in other tolerance mechanisms, such as oxidative stress response signalling [45]. In eggplant, high Pro levels under water stress conditions are associated with lines with higher stress tolerance [37,46]. Thus, Pro accumulation may serve as a useful biochemical marker in the selection of drought-resilient eggplant lines in breeding programs [47].

Chl and Flav pigments were positively correlated with biomass and negatively correlated with water content. This negative correlation is often due to the dilution of pigments in more hydrated tissues, which reduces their apparent concentration [8,48]. Higher chlorophyll content facilitates greater plant growth, but under stress conditions, plants tend to reduce their chlorophyll content [49,50]. However, some more tolerant lines remain unaffected in this regard [51], and in eggplant, chlorophyll content was found to be positively correlated with biomass under water stress [37]. Within the abiotic stress response, flavonols play an important role as antioxidants [52]. Other studies in eggplant suggest that flavonoids are among the principal compounds responding to oxidative stress [11,12].

In this study, three genomic regions were found to be associated with traits relevant to water deficit tolerance. One candidate gene was identified in the genomic region on chromosome 12 associated with growth (Total DW) under stress conditions, out of 31 genes in the region. This gene, *DOF2.1* (SMEL_012g394910.1), which is also associated with stress-induced growth, encodes a DOF (DNA binding with One Finger) transcription factor. These genes are involved in mechanisms such as osmolyte accumulation, photosynthesis and root growth under abiotic stress conditions in Arabidopsis and tomato [53,54]. In potatoes, the expression of these genes has been shown to improve tolerance to water stress [55].

A genomic region on chromosome 11 was associated with Total WC under stress conditions, with four candidate genes out of 41 analysed showing a function related to the trait. The gene *INT4* (SMEL_011g363040.1), which encodes an inositol transporter, may play an important role in stress tolerance, where inositol has been implicated in the regulation of intracellular signalling and auxin regulation that are critical for stress response in Arabidopsis [56]; in pepper, inositol application has been shown to increase water potential and water content in plants under water stress [57]. The gene *Os03g0210100* (SMEL_011g363100.1) encodes a cysteine proteinase inhibitor that promotes resistance to various abiotic stresses, such as drought in Arabidopsis [58]. Regarding the gene *COP1* (SMEL_011g363370.1), an E3 ubiquitin-protein ligase, similar genes in Arabidopsis and pea were found to be involved in the response to term dehydration by stomatal closure through their E3 ubiquitin ligase activity [59]. While the gene *DSP1* (SMEL_011g363380.1), a tyrosine-protein phosphatase, may have a role as a negative regulator of water stress, a similar gene in rice was found to alter the ABA response to stomatal closure, affecting water loss [60]

In the genomic region related to Flav, on chromosome 11, 45 genes were found, of which only one had a described related function. The gene *BHLH75* (SMEL_011g373070.1) encodes a bHLH transcription factor, which has been found to significantly increase flavonoid synthesis in Arabidopsis and tomato and improve the response to water stress by reducing oxidative stress indicators such as hydrogen peroxide and malondialdehyde contents [61,62].

To our knowledge, there are no studies on genomic prediction and selection specifically in eggplant, although there are references to its successful use in other cultivated species of the genus *Solanum* [63–65]. The analysis of the traits evaluated under stress conditions allowed the selection of traits conferring higher tolerance to the lines to generate a selection index that allowed sorting from lines with higher tolerance to those more susceptible. Lines superior to the parents, with high breeding value for stress tolerance, were selected. The PCA groups tolerant and susceptible lines differently, confirming a contrasting phenotypic profile. The *S. melongena* MAGIC parents were found to be within the range of values of the lines. While the parents with the highest tolerance were H15 and AN-S-26, which also showed a good water stress tolerance in previous trials [11], they are also the parents with the highest genetic relatedness [28], the large number of MAGIC lines with a tolerance index above these two parents indicates the appearance of new genetic combinations among MAGIC lines with high breeding potential.

For the traits evaluated in the trial, high variation was found for heritability, genetic variance and predictive ability. While the traits with the highest genetic and phenotypic variation were related to the stem, which is not considered a key indicator of drought tolerance, other traits more commonly associated with water stress tolerance, such as Root growth [43] and Pro [66], exhibited considerable genetic variation coefficients. High genetic variability in the evaluated traits is essential for plant breeding. In this context, broad-sense and narrow-sense heritability values provide insights into the extent to which the observed variation within the population is attributable to total genetic factors or additive genetic factors, respectively [67]. In the MAGIC population, some traits displayed high heritability values (>0.70), whereas others exhibited lower values (<0.30). Traits with low heritability can benefit significantly from genomic selection, where it has been shown to achieve substantial genetic gains [64].

The calculated predictive accuracy ranged from negative to intermediate values. Negative values or values close to zero indicate characteristics the model cannot predict due to low additive variance, while values approaching 0.4 reflect a good predictive ability [31]. The high variability found in the present results has been observed in other studies with polygenic traits [68]. Both forms of cross-validation yield similar predictive results for different traits in our study, and these methods are widely used in plant breeding to evaluate model performance [69,70]. LOO cross-validation uses a larger number of lines for prediction, but it was slightly superior to 5-fold in a smaller number of traits. Similar results were found by Ravelombola et al. (2021) when evaluating a cowpea MAGIC population under water stress conditions [34]. They found traits with medium and low predictive accuracy; however, unlike the results of this study, the larger the training population size, the better the predictions. On the other hand, a predictive ability of up to 0.60 has been demonstrated for complex traits such as yield in a barley MAGIC population [33]. The traits used to construct the selection index showed a prediction accuracy above 0.30 for most of the traits, with proline standing out with a fairly good prediction value; however, the prediction of Flav and Total WC was low. Overall, the results show that the selection of non-phenotyped lines can be moderately well predicted using the model.

This study provides an integrative framework for understanding and improving drought tolerance in eggplant through phenotyping, GWAS, and genomic selection. The first eggplant MAGIC population showed a high phenotypic and genetic diversity, with lines displaying values transgressing the parents for traits such as root biomass and Pro, key traits associated with stress tolerance. In addition, GWAS analysis identified three genomic regions with key genes associated with Total DW, Total WC and Flav, which are associated with greater tolerance to water stress. The application of genomic selection models with index allowed a more accurate selection of tolerant lines with a higher biomass growth, root biomass, flavonol content, water content and proline content. In addition, the predictive accuracy of the model resulted in high values for some traits; however, it did not prove useful for others. In summary, the results obtained demonstrate the potential value of the eggplant MAGIC population for eggplant breeding targeting improved tolerance to water stress.

## Material and Methods

### Plant Material

We evaluated a subset of 184 lines from a MAGIC population of eggplant, along with its eight parental lines. Among the eight MAGIC population parents, seven belong to cultivated eggplant (*S. melongena*) (MM1597, DH ECAVI, AN-S-26, H15, A0416, IVIA-371, and ASI-S-1), whereas the remaining one corresponds to the wild relative *S. incanum* (MM577) [28]. This population, the first of its kind, is known as the MEGGIC (*MAGIC EGGplant InCanum*) population and consists of 325 S5 inbred lines [25]. The 184 lines selected for evaluation were chosen to largely sample the observed genetic and phenotypic variation using the Core Hunter 3 software [71]. Genomic data from 293,783 high-quality SNP markers, along with six phenotypic traits, were employed for line selection to capture the full diversity of the entire population [25]. The SNPs markers were generated through low-coverage whole-genome sequencing (lcWGS) at 3X following Baraja-Fonseca et al. (2024) recommendations [72]. The morphological descriptors included two related to chlorophyll pigmentation in fruit peel, three related to anthocyanin pigmentation (in vegetative plant tissues, fruit peel, and light-insensitive pigmentation under the calyx), and one related to the presence of prickles. These descriptors, combined with the available genomic markers, facilitated the identification of lines with diversity for genetic backgrounds and key agronomic traits, ensuring a robust and representative subset for further evaluation.

Biallelic SNPs from lcWGS (3X) were identified using Freebayes following Baraja-Fonseca et al. (2024) recommendations, benchmarked against a high-confidence reference panel and filtered to remove highly heterozygous, low-depth, monomorphic, and missing data sites [72]. The final SNP dataset was imputed, retaining only markers with MAF > 0.04 and a minimum distance of 2,000 bp, resulting in 293,783 SNPs for linear model analysis and GWAS. These SNPs were obtained from the genotyping data described by Arrones et al. (2025), ensuring the consistency and reliability of the markers used for the analysis (25).

### Growing Conditions

Seeds were germinated in Petri dishes following the Ranil et al. (2015) protocol [73]. After germination, the seeds were placed in seedling trays with a growing substrate (Humin substrate N3, Klasmann-Deilmann, Germany) in a growth chamber. Then, 40 days after sowing, they were transferred to a greenhouse under controlled conditions (maximum 30°C and minimum 15°C) and transplanted into 1.5 L pots with the same growing substrate and fertilised with 7 g per pot of complex fertiliser (Entec® Nitrofoska®, EuroChem, Switzerland) which has a macroelements composition of 14% N, 7% P_2_O_5_, and 17% K_2_O.

The plants were evaluated in three experiments with a completely randomised design. Each experiment represented a separate batch corresponding to a different sowing date, and the experiments were established sequentially (that is, the seeds of the second experiment were sown after the first experiment was finished, and the third after the second). Parental lines were replicated three times per block, while a single replicate within each experiment represented each MAGIC line. Additionally, three extra replicates of the parents and one extra replicate of each of the MAGIC lines were included per experiment to destructively assess the biomass of each parent and line at the start of the water stress treatment. Each replicate consisted of an individual plant.

Plants were watered to 100% field capacity (FC) three times per week for 14 days and then subjected to water stress for 21 days by watering the plants to 30% FC three times per week. The growing substrate FC ranged from approximately 20% before watering (where some plants started to lose turgor) to 30% FC after watering (Figure 6). Watering rates were calculated using the gravimetric method [48], where pots were filled with substrate equivalent to 182 g dry weight, then watered to saturation, covered on the top to avoid evaporation losses, and allowed to drain for approximately 48 h until a constant weight was reached, which was considered the weight of the pot at 100% FC. The amount of water at 100% FC (780 g) and 30% FC (234 g) was obtained by subtracting the weight of the pot and the dry substrate.

**Figure 6.**
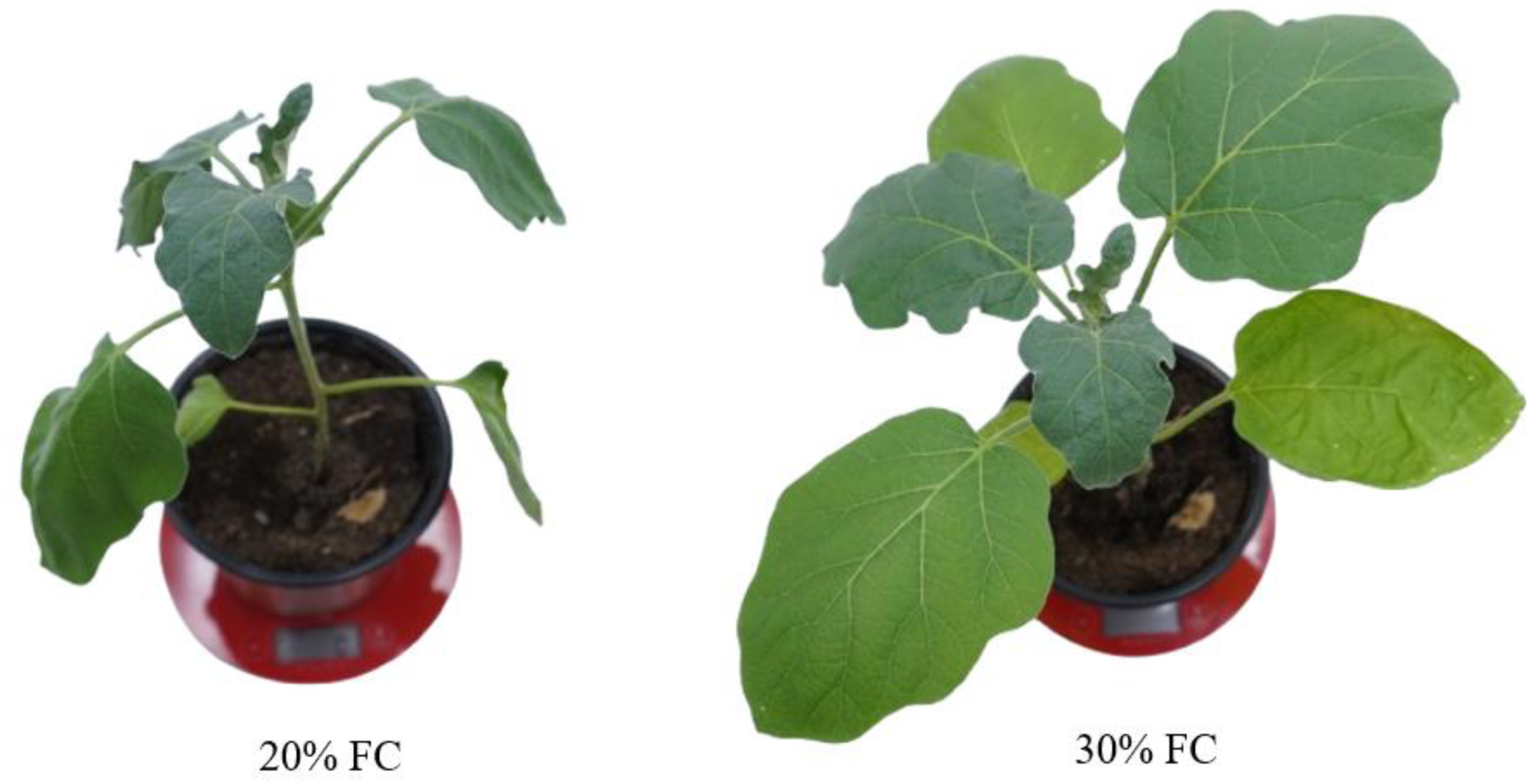
Representative plant of an eggplant MAGIC line during the water stress treatment with a growing substrate water content corresponding to 20% (before irrigation) and 30% (after irrigation) of the field capacity (FC).

### Phenotyping

Before starting the water stress treatment and after 21 days of water stress, the number of leaves was counted, and the stem length was measured. A destructive analysis was also performed at the start of the water stress on one plant from each line in each experiment. This allowed the determination of final values for each trait and also the increases during the period of water stress. The fresh and dry weights of each organ were used to calculate the WC for each plant, and the DW was used to calculate the aerial to root weight ratio.

One day before the end of the experiments (i.e., 20 days after starting the irrigation reduction treatment), Chl, Flav, Anth and NBI were determined using a Dualex® optical sensor (Force-A, Orsay, France). Data were collected from the adaxial and abaxial sides of two developed leaves from the top of each plant.

Pro content was measured spectrophotometrically according to the protocol of Bates et al. (1973) [74]. To extract Pro from leaf samples, 0.1 g was ground, and 1 mL of 3% sulphosalicylic acid was added. The extracts were centrifuged, and 0.5 mL of the supernatant was taken and mixed with 0.5 mL of ninhydrin acid and 0.5 mL of glacial acetic acid. The samples were incubated at 96°C for one hour, and then three mL of toluene was added. The absorbance was measured at 520 nm, and the concentration was quantified using an L-proline standard curve. Pro contents were expressed as (µmol g^-1^ DW)

### Statistical Analysis

To compare the means and ranges of values for each trait, adjusted means were calculated using a linear model specified as:

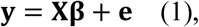

where **y** is the vector of phenotypic observations across 3 experiments with *n* plants per experiment, **β** is the vector of fixed effect estimates for lines and experiments, with associated design matrix **X**, and **e** is the vector of the residuals, which are assumed to be normally distributed as **e** ∼ **N**(0, σ_e_^2^ **I**), where **I** is the identity matrix.

To predict the total genetic effects of MAGIC lines in the experiment for each evaluated trait, genomic values were predicted using a linear mixed model fitted using *sommer* package in R [75], specified as:

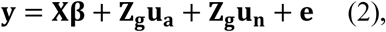

where **y** is the vector of phenotypic observations and **β** is the vector of fixed effect estimates for lines and experiments, with associated design matrix **X**. The total genetic effects of the lines were partitioned into two components: additive and residual genetic effects. The vector of additive genetic effects, **u_a_** ∼ **N**(0, σ_a_^2^ **G**), represents the genomic estimated breeding values, with associated design matrix **Z**_g_, where **G** is the genomic relationship matrix constructed using the method of Van Raden (Van Raden 2009). The vector of residual genetic effects, **u**_n_ ∼ **N**(0, σ_n_ ^2^ **I**), with associated design matrix **Z**_g_ capture genetic variation not accounted by **G** and are modelled using identity matrix **I**. Finally, **e** is the vector of residuals, which are assumed to be normally distributed as **e** ∼ **N** (0, σ_e_^2^ **I**).

Broad-sense (H^2^) and narrow-sense (h^2^) heritabilities were calculated from the model in Equation 2 following Cullis et al. (2006) [76]:

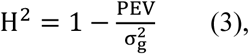

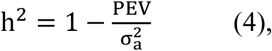

where σ_g_^2^ and σ_a_^2^ are the total and additive genetic variances, and PEV is the mean prediction error variance of genotypic estimates. In addition, the genetic coefficient of variation was calculated as gCV = σ_g_^2^/μ for each trait to compare genetic variation across traits at the genetic level.

To examine the relationship between traits and lines, a principal component analysis (PCA) was conducted on standardised predicted genetic values from the model in Equation 2 using the *stats* package. Pearson correlations of MAGIC lines with the genetic values were computed using the *psych* package [77]. All multivariate analyses were performed in R (R Core Team, 2024).

### Cross-validation and genomic prediction

Two cross-validation strategies were used to assess how well the model in Equation 2 could predict the additive genetic values of MAGIC lines that were not included in the experiment. The first strategy was the leave-one-out cross-validation (LOOCV), where each line is iteratively left out of the training set and predicted using the rest. The second was a 5-fold cross-validation, where the dataset is randomly divided into five subsets, with each subset used once as a validation set while the others form the training set. The 5-fold cross-validation was repeated 100 times. Predictive ability was calculated as the Pearson correlation coefficient (r) between the adjusted means from the model in Equation 1 and predicted additive genetic values in the validation set. Prediction accuracy was calculated by adjusting predictive ability by the square root of the h^2^ calculated from the model in Equation 2 with complete phenotypic data [78]. Once the prediction accuracies had been calculated for all traits, Equation 2 was used to predict the additive genetic values of the remaining 141 MAGIC lines that had not been phenotyped.

### Genome-wide association study and candidate genes

The genome-wide association study (GWAS) was performed using adjusted means from the model in Equation 1, and the analysis was conducted using a general linear model (GLM) in Trait Analysis by aSSociation, Evolution, and Linkage (TASSEL) software (ver. 5.2, Bradbury et al. (2007) [79]). The results were visualised using the R package *qqman* [80], and SNP significance was determined based on a Bonferroni threshold, calculated as: –log_10_ α / total number of SNPs, where α = 0.05.

Only significant SNPs, along with several nearby markers showing high –log₁₀(*p*) values (greater than 5), were considered to determine the genomic region. Linkage disequilibrium was then calculated using all SNPs in the region to delimit the associated genomic interval. Candidate genes were identified within this region using the Sol Genomics Network database [81].

### Index-based selection of drought-tolerant lines

In order to identify lines with a higher tolerance to drought stress, a genomic selection index was constructed based on six drought-related traits: Total DW, Total DW growth, Root DW, Flav, Total WC and Pro. These traits represent different clusters in the correlation analysis. Total DW and Total DW growth cluster with Pro, while Flav clusters with Root DW, and Total WC forms a separate cluster. These traits were selected because of their direct relevance to the plant’s ability to cope with water stress. Total DW is a crucial trait as it ensures biomass gains under stress, indicating overall plant growth and productivity, even in conditions of limited water availability [82]. Root DW is vital for assessing the plant’s root development, as it allows greater water absorption [83]. Flav plays an important role in protecting plants from oxidative stress [84]. Total WC is a critical indicator of plant hydration, helping to select plants that maintain adequate water levels under drought conditions [85]. Lastly, Pro serves as an osmoregulatory compound, helping the plant to retain water during periods of dehydration, while also functioning as an antioxidant and playing a role in stress signaling pathways, further enhancing drought tolerance [45].

The selection index values were calculated using the following formula:

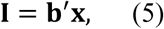

where **x** is a vector of additive genetic effects for each trait obtained from the model in Equation 2, and **b** is a vector of index coefficients calculated using the Smith-Hazel formula **b** = **P^−1^Ga** [86]. Phenotypic and genetic covariance matrices, **P** and **G**, were calculated as the covariances between genetic values and phenotypic values, respectively, and the vector of economic weights **a** was set to 1 for all five traits. Lines showing the highest index values were classified as tolerant and lines with the lowest index values classified as susceptible. Index values were also calculated for untested predicted lines to identify potentially tolerant or susceptible lines.

## Acknowledgments

This work was funded by MICIU/AEI/10.13039/501100011033 and by ERDF/EU (PID2024-160953OB-I00) and co-funded by the European Union. Funding was also received from grant CIPROM/2021/020 funded by Conselleria d’Educació, Universitats i Ocupació of the Generalitat Valenciana, and from grant PDC2022-133513-I00 funded by MICIU/AEI/10.13039/501100011033 and by the European Union NextGeneration EU/PRTR. MF-S is grateful to Conselleria d’Educació, Universitats i Ocupació of the Generalitat Valenciana for a pre-doctoral grant within the Santiago Grisolía program (GRISOLIAP/2021/151). AA is grateful to Conselleria d’Educació, Cultura, Universitats i Ocupació of the Generalitat Valenciana and to the FSE+ (European Social Fund Plus) of the European Union for a post-doctoral contract within the CIAPOST program (CIAPOS/2024/330). PG has received a postdoctoral grant (RYC2021-031999-I) funded by MICIU/AEI/10.13039/501100011033 and by the European Union NextGeneration EU/PRTR. Funding for open access: Universitat Politècnica de València.

## Supplementary data

**Fig. S1.**
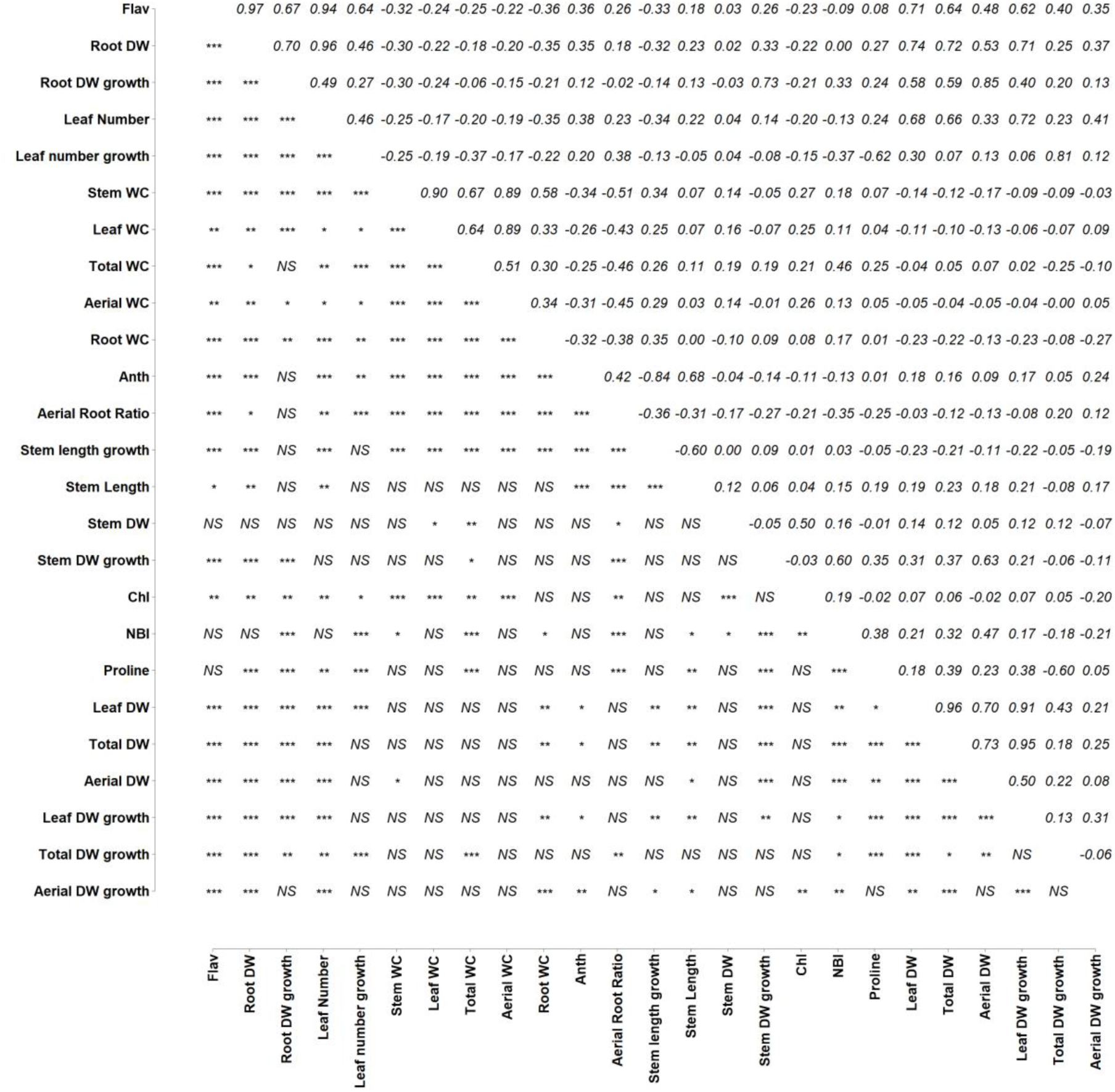
Correlation matrix coefficients in a subset of 184 MAGIC lines. Upper right diagonal shows the correlation coefficient and lower diagonal shows the significant correlation. ^ns^, *, **, *** indicate non-significant for a p-value < 0.05 and significant for a p-value < 0.05, < 0.01 and < 0.001, respectively.

**Fig. S2.**
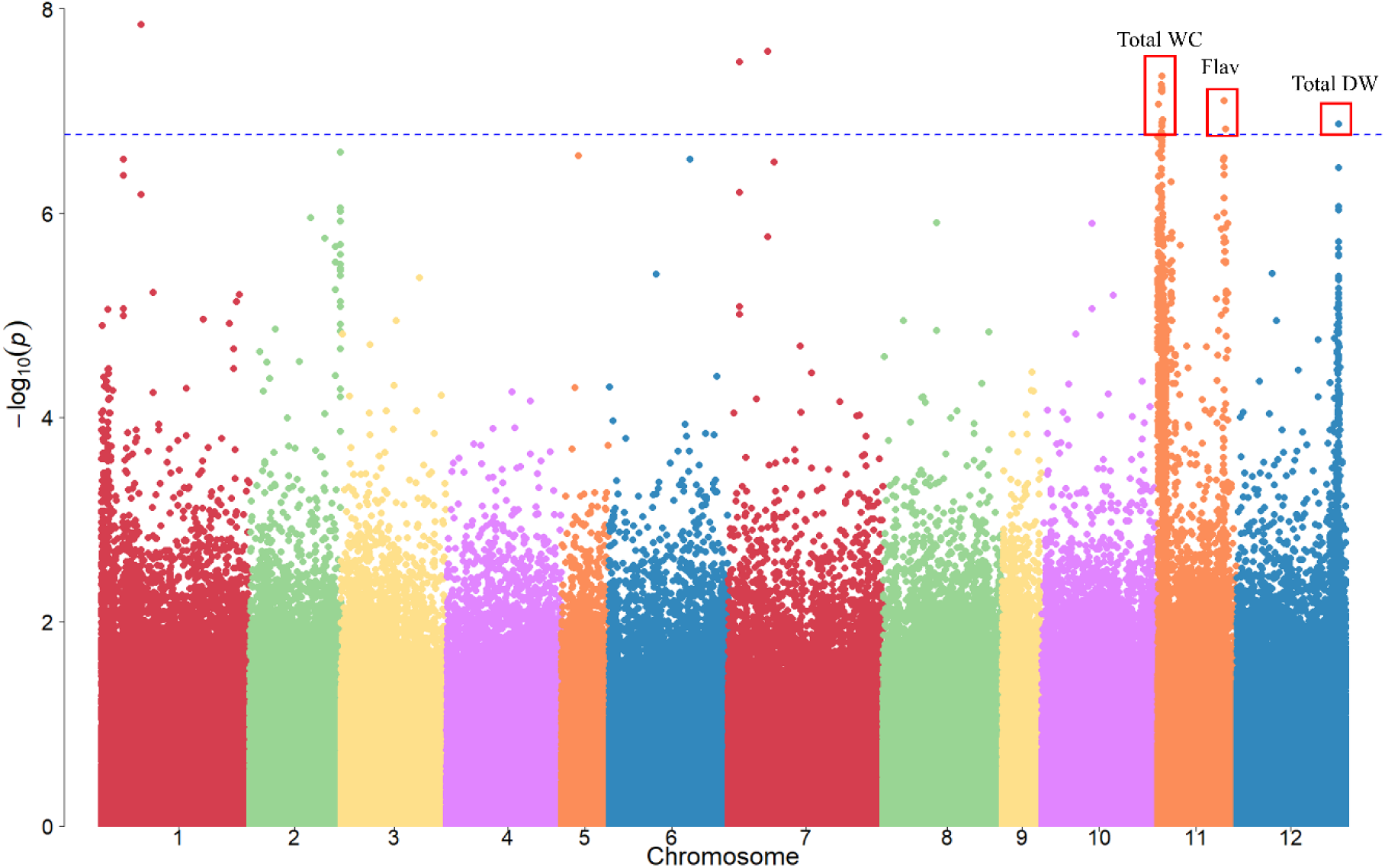
Manhattan plot of the traits Total WC (water content), Flav (flavonol index) and Total DW (dry weight) in the GWAS. The x-axis represents the 12 eggplant chromosomes. The y-axis represents the –log₁₀(*p*) values. Significant SNPs defining a genomic region for a trait are marked in red.

